# Predicting bacterial fitness in Mycobacterium tuberculosis with transcriptional regulatory network-informed interpretable machine learning

**DOI:** 10.1101/2024.09.23.614645

**Authors:** Ethan Bustad, Edson Petry, Oliver Gu, Braden T. Griebel, Tige R. Rustad, David R. Sherman, Jason H. Yang, Shuyi Ma

## Abstract

*Mycobacterium tuberculosis* (Mtb) is the causative agent of tuberculosis disease, the greatest source of global mortality by a bacterial pathogen. Mtb adapts and responds to diverse stresses such as antibiotics by inducing transcriptional stress-response regulatory programs. Understanding how and when these mycobacterial regulatory programs are activated could enable novel treatment strategies for potentiating the efficacy of new and existing drugs. Here we sought to define and analyze Mtb regulatory programs that modulate bacterial fitness. We assembled a large Mtb RNA expression compendium and applied these to infer a comprehensive Mtb transcriptional regulatory network and compute condition-specific transcription factor activity profiles. We utilized transcriptomic and functional genomics data to train an interpretable machine learning model that can predict Mtb fitness from transcription factor activity profiles. We demonstrated that this transcription factor activity-based model can successfully predict Mtb growth arrest and growth resumption under hypoxia and reaeration using only RNA-seq expression data as a starting point. These integrative network modeling and machine learning analyses thus enable the prediction of mycobacterial fitness under different environmental and genetic contexts. We envision these models can potentially inform the future design of prognostic assays and therapeutic intervention that can cripple Mtb growth and survival to cure tuberculosis disease.

## 1. Introduction

*Mycobacterium tuberculosis* (Mtb) remains a supremely successful pathogen, sickening 10.6 million people and killing over 1 million people worldwide each year [1]. An important factor for Mtb’s success is its ability to adapt to a broad range of host-associated and treatment-associated stresses. The mechanisms underlying how Mtb dynamically regulates its growth and physiology in response to stress response remains incompletely understood. Characterizing the gene regulatory activities of transcription factors (TFs) under different environmental or stress conditions could help inform interventions that modulate Mtb growth and survival to cure tuberculosis disease.

Several groups have previously performed analyses to characterize Mtb’s transcriptional regulatory network (TRN) using experimental and computational approaches [2; 3; 4; 5; 6; 7; 8; 9]. These efforts have largely relied on two strategies: 1) detailed profiling of the molecular impact of individual transcription factors (TFs) with recombinant induction and disruption strains, and/or 2) statistically informed TRN inference using data from large transcriptome compendia.

In principle, TRNs can be empirically assembled from measurements of TF-DNA binding activities and gene expression profiles from conditions with known individual TF perturbations. These data would enable the inference of direct regulatory interactions between TFs and their putative target genes, which exhibit altered transcriptional expression in response to TF perturbations and provide evidence of TF binding events proximal to a gene. To leverage this strategy, we previously engineered a library of Mtb recombinant TF induction (TFI) strains [2; 6], from which we profiled transcriptomes in 208 TFI strains by microarray analyses (GSE59086, [6; 10]) and detected ∼16,000 ChIP-seq binding events for 154 TFs (∼80% of all Mtb TFs) and 2,843 genes (∼70% of all Mtb genes) [3; 10]. These detailed ChIP-seq and transcriptional profiles have yielded important insights into the regulatory programs active during Mtb broth culture. However, these experiments possessed several technical limitations. For example, our microarray profiling efforts were unable to measure changes in expression for 1,190 genes (∼30% of Mtb genes) [6], and our ChIP-seq profiling efforts were unable to detect TF binding associated with 1,040 genes (∼26% of Mtb genes) [3].

Moreover, the existing profiles have focused specifically on regulatory behavior of the Mtb laboratory strain H37Rv in log-phase growth in 7H9 media. Consequently, condition-specific interactions relevant to other environments or Mtb strains were not captured. Thus, despite such efforts, significant gaps remain in the ability to identify TF-gene regulatory interactions directly and comprehensively by only experimental activities.

Bioinformatic network inference approaches that utilize expression compendia comprising transcriptome responses under diverse biological conditions are a useful complementary strategy to recombinant strain profiling. These statistically informed approaches enable assessment of regulatory interactions across the multitude of conditions present in a transcriptome compendium. However, these computational network inference strategies are constrained by two limitations. First, large and biologically diverse gene expression data are needed to fuel identification of high-confidence statistical associations between TFs and putative target genes [11]. To meet this need, compendia of expression data may be curated from public microarray [4; 10] or RNA-seq [7; 12; 13] data. Second, statistical learning network inference algorithms differ in the assumptions made on the training data and on the interpretation of TF-gene associations. These assumptions are often biologically inaccurate. We previously performed such analyses and were able to only infer 598 clusters of coregulated gene expression for 3,922 genes [4]. Others recently performed similar analyses and inferred either 80 clusters for 3,906 genes [7] or 560 co-regulated gene modules for 3,912 genes [5]. These models have successfully revealed novel regulatory interactions impacting Mtb stress adaptation, but none of these regulatory models may be precisely interpreted as TF regulatory programs (as they only capture a fraction of Mtb’s 214 TFs) and none can be used to directly estimate TF activities (i.e., the extent of regulation that each TF exerts on its regulated target genes, TFAs, [14]) under different experimental conditions. TRN inference efforts in other microbes, including the DREAM5 challenge for *E. coli* and *S. aureus* [15], have found that robust TRNs may be assembled by aggregating the regulatory relationships inferred by different statistical algorithms. We hypothesized that implementing a similar “wisdom of crowds” approach to aggregate complementary TRNs inferred via different statistical approaches would yield a more comprehensive and higher quality Mtb TRN.

Here we assembled a biologically diverse and batch corrected Mtb RNA-seq gene expression compendium. We integrated this RNA-seq compendium with the perturbative TFI microarray dataset to infer a comprehensive Mtb transcriptional regulatory network that included all 214 TFs and all 4,027 genes present in our RNA-seq expression compendium. We used this TRN to estimate TFA profiles corresponding to individual RNA expression profiles. We used the TFAs calculated from our RNA-seq compendium to train an interpretable machine learning regression model that could predict growth phenotypes previously measured in TF-induced strains [16]. We demonstrated that this regression model can accurately predict Mtb fitness under stressful environmental conditions such as hypoxia.

## 2. Methods

### 2.1 TFI microarray expression compendium assembly and normalization

Microarray expression data corresponding to TFI strains were downloaded from GEO (GSE59086). Groups were assigned to each sample by the identity of each strain. The Rv2160A gene fully encompasses the Rv2160c gene, so the Rv2160A and Rv2160c samples were combined into a single Rv2160 TFI strain group. This resulted in 208 TFI strain groups. These 208 strain groups included Rv0560, Rv3164c, and Rv3692 which were considered hypothetical TFs in TFI strain construction [6], but later determined to not be true Mtb TFs [10]. However, for the purpose of the analyses presented here, each of these 208 strains will be referred to as TFs. Smooth quantile normalization [17] was performed using *PySNAIL* [18] using the assigned group definitions.

### 2.2 RNA-seq expression compendium assembly, quality control, and normalization

The NCBI Sequence Read Archive (SRA) was queried with “*Mycobacterium tuberculosis*” for RNA expression samples containing raw FASTQ sequencing reads. 3,506 FASTQ sequencing reads were downloaded and combined with FASTQ sequencing reads from 398 unpublished RNA-seq profiles generated by our labs. We aligned these sequencing reads against the NC_000962.3 Mtb H37Rv reference genome using Bowtie 2 [19]. Read counts were compiled using *featureCounts* [20]. Samples with fewer than 400,000 total gene counts and samples duplicated in our preliminary compendium were excluded from further analysis. Sequencing counts between samples were normalized by transcripts per kilobase million (TPM). Group definitions were manually added to represent unique experimental conditions from each set of experiments; biological replicates for each experimental condition were given the same group definitions. Smooth quantile normalization [17] was performed using *PySNAIL* [18] using the assigned group definitions. Quality data, adapter and quality trimming statistics, and alignment and counts metrics were compiled and assessed using *MultiQC* [21].

### 2.3 UMAP visualization and cluster estimation

RNA expression compendia and TFAs were visualized by Uniform Manifold Approximation & Projection (UMAP) [22]. Clusters were estimated by *DBSCAN* [23]. The ε hyperparameter was optimized for each dataset by varying ε across 50 logarithmically distributed values from 0.1 to 10 and selecting the value of the elbow of the ε vs. Number of Outliers plot. This selection delivers the minimum number of clusters that maximizes inclusion of samples without overfitting the data (**Supplementary Figure S1**). UMAP and DBSCAN analyses were performed in Python using their implementations in *umap-learn* and *scikit-learn* [24].

### 2.4 Regulatory network inference methods

We implemented an ensemble of network inference methods by starting with a selection of methods featured in the DREAM5 challenge [15]. These methods were selected based on diversity in underlying statistical approach, predictive performance reported in the DREAM5 study, and the availability of a working implementation. Our initial selection consisted of ARACNe [25; 26], CLR [27], and GENIE3 [28]. We chose an ARACNe implementation that employs adaptive partitioning for more efficient processing [25; 26]. We used an R implementation of CLR available on CRAN from the *parmigene* package [29]. We used an R implementation of GENIE3 available on BioConductor [30]. To supplement these methods, we incorporated two other more recent advances in network inference approaches: cMonkey2 [31; 32] and iModulon [33]. We used a docker image containing a Python implementation of cMonkey2, available at https://hub.docker.com/r/weiju/cmonkey2. For iModulon, our desired output was different from the output of this algorithm implemented by the original authors. We thus made a custom implementation, borrowing heavily from https://github.com/SBRG/pymodulon and https://github.com/SBRG/iModulonMiner, in Python. In addition, we also chose to implement a regression strategy using Elastic Net regression, a more advanced technique than was used in DREAM5. Elastic Net is a regularization method that takes advantage of the unique properties of both the lasso (used extensively in DREAM5) and ridge regression [34]. Elastic Net performs better than lasso or ridge regression when predictors may be correlated and under-determined [35]. We modeled each gene individually on the expression of all the transcription factors, and used the resulting coefficients to both select significant relationships and score those relationships; this implementation was done in Python using *scikit-learn* [24]. Descriptions of each of these inference methods are provided in **Supplementary Table 3**.

Each method was wrapped to produce a ranked list of putative TF regulator-target gene relationships in order of the inferred strength of the regulatory relationship, from strongest to weakest. Execution was done using docker images (https://hub.docker.com/repositories/malabcgidr?search=network-inference). Auto-regulatory (self-targeting) relationships were excluded. Method hyperparameters were chosen to match either original publications or the DREAM5 challenge when possible. Execution for each method and optimization of their corresponding hyperparameters was validated by testing against the evaluation scripts provided in the supplemental material of [15; 32].

A network was generated for each combination of the two datasets (RNA-seq and TFI microarray) and 6 inference methods, yielding 12 total constituent networks.

### 2.5 Inferred network truncation and aggregation

The constituent networks were large, as many of the network inference methods did not require a cutoff threshold and did not perform multiple testing correction; the union of all inferred edges constituted over 90% of the possible Mtb regulatory space (where 100% would be every TF harboring a regulatory association with every Mtb gene). We therefore truncated each inferred network to incorporate the unique perspective of each model without aggregating too many low-confidence relationships. This was done by comparison with an independent validation set, comprising a presumed unbiased sampling of the true population of regulatory relationships in Mtb. This validation set was used to identify the extent of true positives in each network.

The validation data set was gleaned from Sanz et al., Material S1 [8]. The original list was filtered for relationships whose supporting evidence included at least one high-confidence physical methodology, namely values 4-9: LacZ-promoter fusion, GFP-promoter fusion, proteomic studies, electrophoretic mobility shift assays (EMSA), one hybrid reporter system, and chip-on-chip. This yielded a set of 433 high-confidence regulator-target relationships, including 51 regulators and 160 total target genes, that had little to no dependence on the transcriptional information used to build the constituent networks.

A cutoff threshold was chosen for each network by binning the ranks of validation hits into 32 bins and truncating the network at the first bin where the number of hits fell below the expected level of random overlap per bin. This level was calculated to equal the mean of a hypergeometric distribution, with a population size equal to the total regulatory space of Mtb, a set of true positive regulatory interactions identified by the Sanz validation set [8], and draws equal to the size of the inferred network, taken without replacement. This shrunk each network to an average of about 10% of its original size (3-28%) (**Supplementary** Figure 2). Three of the constituent networks displayed insufficient enrichment against the validation dataset: ARACNe/TFI, cMonkey2/TFI, and iModulon/TFI. Upon executing a Fisher’s exact test to determine the chance of a random network achieving the same enrichment, these three failed to pass a strict cutoff of 0.0001. They were thus excluded from further aggregation.

The remaining truncated networks were then aggregated together, first into two combined networks, one for each underlying input transcriptome dataset (RNA-seq compendium and TFI microarray profile). Aggregation was performed by rank average as described in the DREAM5 challenge [15]. Repeating the enrichment analysis performed above, it was determined that the TFI aggregate would benefit from additional truncation and was thus truncated using the same threshold strategy described in the previous paragraph, whereas the RNA-seq network was already sufficiently enriched. These two networks were then aggregated together again by rank average, yielding one final aggregate network.

All these networks were validated against the Sanz et al. data set using the Matthews Correlation Coefficient (MCC), as described previously [36; 37] (**Supplementary** Figure 3).

### 2.6 Principal Component Analysis

Principal component analysis (PCA) was performed on the inferred networks (after truncation), the dataset-level aggregate networks, and the overall aggregate network, using the 16,792-dimensional space represented by the ranks of edges shared across at least 3 of the inferred networks. Any relevant edges not included in a given network were assigned a rank of 16,792, the size of the space.

### 2.7 Regulatory directionality

The types of the regulatory connections (whether the TF up- or down-regulates the associated gene) were explored using a combination of the regression models and measured TFI gene expression values. Two elastic net models and two unpenalized linear models were used to infer direction of regulation based on the sign of the regression coefficients, one of each for each dataset (RNA-seq compendium and TFI microarray profile). We supplemented these regression associations with the directionality of significant differential gene expression (i.e. upregulated vs. downregulated expression) measured from the TFI microarray dataset. Linear models were fit in Python with the *statsmodels* package. Coefficients with an FDR < 0.05 were selected as evidence. Elastic net models with an R^2^ of less than 0.8 were excluded; coefficients that were included by the remaining models were selected as evidence. TFI differential expression from the microarray dataset was filtered using an FDR < 0.05 and requiring at least 2-fold change in either direction. Elastic net models and TFI differential expression were considered strong evidence, whereas the unpenalized linear models were considered weak evidence. A flow chart depicting how the information from these models and differential expression analyses were used to define up vs. down regulation is shown in **Supplementary** Figure 5.

### 2.8 Comparing inferred networks against independent reference information

Additional orthogonal datasets were incorporated to corroborate the networks. All generated networks were tested against a set of published ChIP-seq binding relationships gleaned from Minch, et al. [3]. We took the intersection of their sets of statistically significant peaks (Supplementary Data 1 from [3]) and peaks in a canonical promoter region (Supplementary Data 3 from [3]) to yield 5,178 relationships, including 129 regulators and 2,271 total targets. The MCC was then calculated against this data set for each network.

Gene ontology enrichment analysis was then performed to ascertain the extent to which TF targeting could be used to gauge biological function within each group [38; 39]. For each TF, each set of genes that our network identified as upregulated, downregulated, or regulated in both directions by the regulator was analyzed for GO enrichment at an FDR < 0.05. All identified GO annotations that had a child annotation also identified for a given TF were removed for the sake of simplicity (**Supplementary Table 5C**). Results were filtered to regulators receiving at least 3 significant GO enrichments for further manual inspection and analysis (**Supplementary Tables 5A, 5B**), and those TFs with an annotated name and considered to have a testably specific functional role listed in the Mycobrowser annotation [40] were juxtaposed for network validation (**Table 1**). GO analysis was performed in Python using the *goatools* package [41]. Gene ontology data was taken from the 2024- 06-17 release of go-basic.obo from the Gene Ontology knowledgebase [42] (https://purl.obolibrary.org/obo/go/releases/2024-06-17/go-basic.obo), and mappings to Mtb genes were taken from the European Bioinformatics Institute GOA project, release 20240805 (https://ftp.ebi.ac.uk/pub/databases/GO/goa/proteomes/30.M_tuberculosis_ATCC_25618.goa).

**Table 1.** Network regulators: annotation versus gene set enrichment analysis of inferred regulon.

| Regulator | Name | Mycobrowser gene product and function information | Inferred Regulon GO Annots. (FDR <0.05) |  |
| --- | --- | --- | --- | --- |
|  |  |  | # | Summary |
| **Rv0353** | hspR | Probable MerR family heat shock protein transcriptional repressor. Involved in repression of heat shock proteins. Binds to three inverted repeats in the promoter region of the DnaK operon. Induced by heat shock. | 3 | heat response |
| **Rv1657** | argR | Probable arginine repressor (AHRC). Regulates arginine biosynthesis genes. | 4 | cobalamin synthesis; UMP synthesis; C-N bond formation |
| **Rv2215** | dlaT | Dihydrolipoamide acyltransferase, component of pyruvate dehydrogenase. Involved in TCA cycle; converts pyruvate to acetyl-CoA and CO<sub>2</sub>. Also involved in defense against oxidative stress. | 51 | TCA cycle, respiration, downregulation of virulence factors |
| **Rv2359** | zur | Probable zinc uptake regulation protein. Acts as a global negative controlling element, with Zn<sup>2+</sup> binds operator of repressed genes. | 8 | downregulating translation, iron import |
| **Rv2374c** | hrcA | Probable heat shock protein transcriptional repressor. Involved in repression of class I heat shock proteins. Prevents heat-shock induction of these operons. | 17 | transcription and translation |
| **Rv2610c** | pimA | Alpha-mannosyltransferase. Involved in the first mannosylation step in phosphatidylinositol mannoside biosynthesis (transfer of mannose residues onto PI). | 64 | amino acid and nucleobase synth., respiration, growth/proliferation |
| **Rv2720** | lexA | Repressor. Represses genes involved in nucleotide excision repair and SOS response. Binds 14-bp palindromic sequence. | 10 | DNA binding, repair, cleavage |
| **Rv3301c** | phoY1 | Probable transcriptional regulatory protein PhoU-homolog 1. Involved in regulation of active transport of inorganic phosphate across the membrane. | 18 | ETC, oxidative phosphorylation |
| **Rv3417c** | groEL1 | 60 kDa chaperonin 1 (protein CPN60-1). Prevents misfolding, promotes refolding and proper assembly of unfolded polypeptides generated under stress conditions. | 15 | stress response |
| **Rv3574** | kstR | Transcriptional regulatory protein (probably TetR-family). Involved in transcriptional mechanism. Predicted to control regulon involved in lipid metabolism. | 22 | cholesterol, lipid, and carbon metabolism |
| **Rv0599c** | vapB27 | Possible antitoxin. | 13 | growth regulation, toxin sequestration, RNase |
| **Rv0608** | vapB28 | Possible antitoxin. | 12 | growth regulation, toxin sequestration, RNase |
| **Rv0623** | vapB30 | Possible antitoxin. | 12 | growth regulation, toxin sequestration, RNase |
| **Rv1560** | vapB11 | Possible antitoxin. | 6 | growth regulation |
| **Rv1740** | vapB34 | Possible antitoxin. | 7 | growth regulation |
| **Rv1960c** | parD1 | Possible antitoxin. | 18 | growth regulation, toxin sequestration, RNase |
| **Rv2009** | vapB15 | Antitoxin. | 13 | growth regulation, RNase |
| **Rv2595** | vapB40 | Possible antitoxin. | 8 | growth regulation, toxin sequestration |
| **Rv2760c** | vapB42 | Possible antitoxin. | 4 | growth regulation, DNA repair |

### 2.9 Calculating transcription factor activity profiles from network component analysis

Transcription factor activities for each expression profile was computed using Robust Network Component Analysis (ROBNCA) [43]. ROBNCA was implemented in Python, using code adapted from https://github.com/CovertLab/WholeCellEcoliRelease/tree/00cf7738cb8379c14d65ef632b2156bdf7c23434/reconstruction/ecoli/scripts/nca [44].

### 2.10 Associating network activity with bacterial fitness

We built a model associating mycobacterial growth with TF activity, as inferred from measured gene expression data. The GSE59086 microarray dataset was again used as a broad measure of TFI conditions, with relative growth data for 194 matching TFI conditions added from Ma et al., 2021, Table S1 as training data [16]. Expression levels in the form of log-2 fold-change were transformed into putative TFAs using the control strengths calculated via NCA from the aggregate network and RNA-seq compendium. A gradient boosted machine (GBM) model was trained to regress growth on TFAs, using a grid search cross-validation scheme to optimize hyperparameters based on bounds derived from [34], using the number of estimators to reward better performing models. The number of estimators was then optimized with a simple grid search. The model was implemented in Python using the *lightgbm* package [45; 46].

### 2.11 Hypoxia time-course experiment

Wildtype H37Rv (ATCC 27294) and H37Rv transformed with a control anhydrotetracycline (ATc)-inducible expression vector (H37Rv::pEXCF-empty, which does not induce recombinant gene expression) were cultured under in Middlebrook 7H9 with the oleic acid, bovine albumin, dextrose, and catalase (OADC) supplement (Difco) and with 0.05% Tween 80 at 37°C. H37Rv::pEXCF-empty was grown with the addition of 50 µg/ml hygromycin B to maintain the plasmid and induced with 100ng/mL ATc one day prior to onset of hypoxia. For hypoxia, strains were cultured in oxygen-limited conditions (1% aerobic O2 tension) for 7 days, followed by reaeration on day 7-12, initiated by transferring cultures into continuously rolled bottles with 5:1 head space ratio using methods described previously [2; 47; 48; 49]. Bacterial survival and growth were enumerated by plating for colony forming units (CFU) on Middlebrook 7H10 solid media plates using standard microbiological methods.

Transcriptomes were generated by RNA-seq from bacterial cultures sampled from the aforementioned conditions using methods described previously [50]. Briefly, bacterial pellets suspended in TRIzol were transferred to a tube containing Lysing Matrix B (QBiogene) and vigorously shaken in a homogenizer. The mixture was centrifuged, and RNA was extracted from the supernatant with chloroform, followed by RNA precipitation by isopropanol and high-salt solution (0.8 M Na citrate, 1.2 M NaCl). Total RNA was purified using a RNeasy kit following the manufacturer’s recommendations (Qiagen). rRNA was depleted from samples using the RiboZero rRNA removal (bacteria) magnetic kit (Illumina Inc., San Diego, CA). Illumina sequencing libraries were prepared from the resulting samples using the NEBNext Ultra RNA Library Prep kit for Illumina (New England Biolabs, Ipswich, MA) according to the manufacturer’s instructions, and using the AMPure XP reagent (Agencourt Bioscience Corporation, Beverly, MA) for size selection and cleanup of adaptor-ligated DNA. We used the NEBNext Multiplex Oligos for Illumina (Dual Index Primers Set 1) to barcode the libraries to enable sample multiplexing per sequencing run. The prepared libraries were quantified using the Kapa quantitative PCR (qPCR) quantification kit and sequenced at the University of Washington Northwest Genomics Center with the Illumina NextSeq 500 High Output v2 kit (Illumina Inc., San Diego, CA). The sequencing run generated an average of 75 million base-pair paired-end raw read counts per library. Read alignment and gene expression estimation was carried out using a custom processing pipeline in R that harnesses the Bowtie 2 utilities [19; 51], which is publicly accessible at https://github.com/robertdouglasmorrison/DuffyTools, and https://github.com/robertdouglasmorrison/DuffyNGS.

Gene expression data were transformed from log-2 fold-change to putative TFAs using the control strengths calculated via NCA above and run through the GBM model to predict relative fitness level of the Mtb culture as it progressed through the hypoxia time-course.

### 2.12 False discovery rate correction

False discovery rate correction was performed using the two-stage Benjamini-Krieger-Yekutieli method [52].

## 3. Results

### 3.1 Generation of a large and biologically diverse Mtb gene expression compendium for TRN inference

Our previous attempts at TRN characterization utilized microarray expression profiles from recombinant TFI strains as perturbative training data (GSE59086, [6]). However, while this dataset enabled detailed characterization of transcriptional regulation of Mtb physiology during log-phase broth culture, it possessed poor biological diversity. UMAP and DBSCAN analyses reveal that expression profiles from these 698 microarray experiments and 208 TFI conditions only yielded 16 clusters of expression profiles (**Figure 1A**). This poor diversity likely arises from the original experimental design for these data, in which each TFI strain was grown to log-phase in albumin-dextrose-catalase (ADC)-supplemented 7H9 media before isolating RNA. UMAP and DBSCAN analyses suggested that this TFI microarray dataset alone would be insufficient for predicting TFAs corresponding to diverse experimental conditions. Moreover, microarray technologies have poor sensitivity and dynamic range for quantifying gene expression [53]. We found that 101 genes in this dataset did not possess expression measurements greater than 10 counts, indicating poor detection or poor evidence for expression in these experiments (**Figure 1B**). In addition, the median absolute deviation (MAD) was small (< 1) for nearly all genes, indicating the ability to detect gene expression changes across conditions was limited. These analyses collectively motivated the need to assemble a new RNA expression compendium.

**Figure 1:**
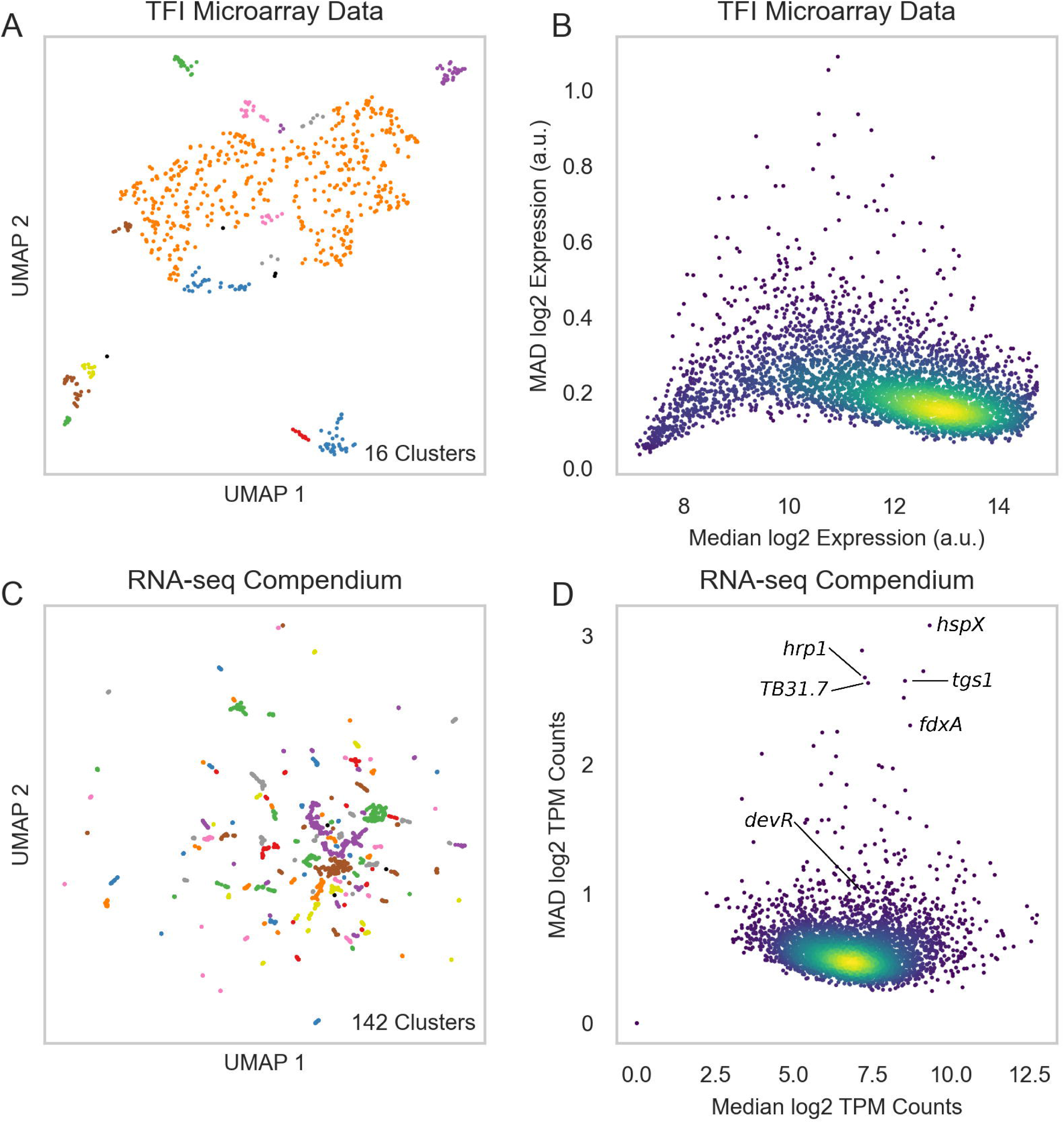
A biologically diverse Mtb RNA expression compendium. (A) UMAP visualization of biological diversity in the TFI microarray data. TFI data were batch corrected by smooth quantile normalization before computing the UMAP. Density-based spatial clustering (DBSCAN) was performed on the UMAP to identify clusters of samples with similar gene expression. UMAP and DBSCAN analyses revealed 16 total expression clusters in the TFI dataset. (B) Median vs. median absolute deviation (MAD) plot of expression for each gene across the TFI dataset. Each point represents a gene. Median expression and MAD were calculated for each gene across the 698 samples. Colors reveal point density (yellow: high density, blue: low density). (C) UMAP visualization of samples from the normalized and batch corrected RNA-seq compendium determined by gene expression. UMAP and DBSCAN analyses reveal 142 clusters of samples with similar gene expression. (D) Median vs MAD plot of expression for each gene across the RNA-seq compendium.

We therefore collected samples from the NCBI Sequence Read Archive (SRA) and our own labs, aligned, filtered, normalized, and batch corrected by smooth quantile normalization [17; 18] (see **Methods** for details). Batch correction is an important pre-processing step for unifying data from different sources that is frequently overlooked in Mtb RNA expression compendium analyses [4; 7; 12; 13]. After performing these pre-processing steps, our final compendium comprised 3,496 RNA-seq samples from 1,288 experimental conditions (**Supplementary Table 1**). Expression counts for the RNA-seq compendium can be queried at https://tfnetwork.streamlit.app/.

UMAP and DBSCAN analyses of the batch corrected RNA-seq expression compendium validated its biological diversity (**Figure 1C**-**D****, Supplementary Table 2**). We identified 142 unique expression clusters. This RNA-seq transcriptome compendium exhibited significantly greater dynamic range and variation in gene expression than in the TFI microarray dataset (**Figure 1D**). Of note, genes with high variation (high MAD) were mostly well-characterized stress response genes (e.g., Rv2031c *(hspX)*, Rv2626c *(hrp1)*, and Rv2623 *(TB31.7)*), with Rv2007c *(fdxA)* having higher variation than the commonly studied Rv3133c *(devR)* stress response regulator. These are consistent with expectation, as most stress response genes would be expected to only be induced in the presence of their specific stressor.

### 3.2 Inferred transcriptional regulatory network interactions enrich for shared functional processes

Network inference studies in other bacteria have shown that combining regulatory interactions from multiple different inference algorithms results in a TRN that outperform networks generated by a single method [15]. To more comprehensively characterize Mtb regulatory interactions, we applied a “wisdom of crowds” ensemble inference approach. We first applied a collection of regulatory network inference tools to generate TRN models using individual methods (see **Methods**). These tools were selected because they have been shown to be sensitive to distinct types of regulatory relationships in other bacteria [15] or they have previously been successfully applied to infer regulatory relationships in Mtb [4; 5; 7]. To further diversify the regulatory relationships inferred from these approaches, we applied these tools to both our assembled RNA-seq compendium as well as the TFI microarray dataset. Collectively, these inference activities yielded 12 networks that describe 779,213 unique interactions between 214 regulators and 4,029 target genes. We truncated these networks using a benchmark dataset of high confidence regulatory interactions with biochemical evidence that was curated by Sanz et al. [8] (see **Methods**). We used this high confidence regulatory interaction dataset to inform pruning of low-confidence regulatory relationships inferred from each of the individual inference methods (**Supplementary** Figure 2), yielding a shorter, more high-confidence network for each method. Principal component analysis of these networks revealed substantial diversity in the regulatory interactions identified between the different approaches applied to the two source datasets (**Figure 2B**).

**Figure 2:**
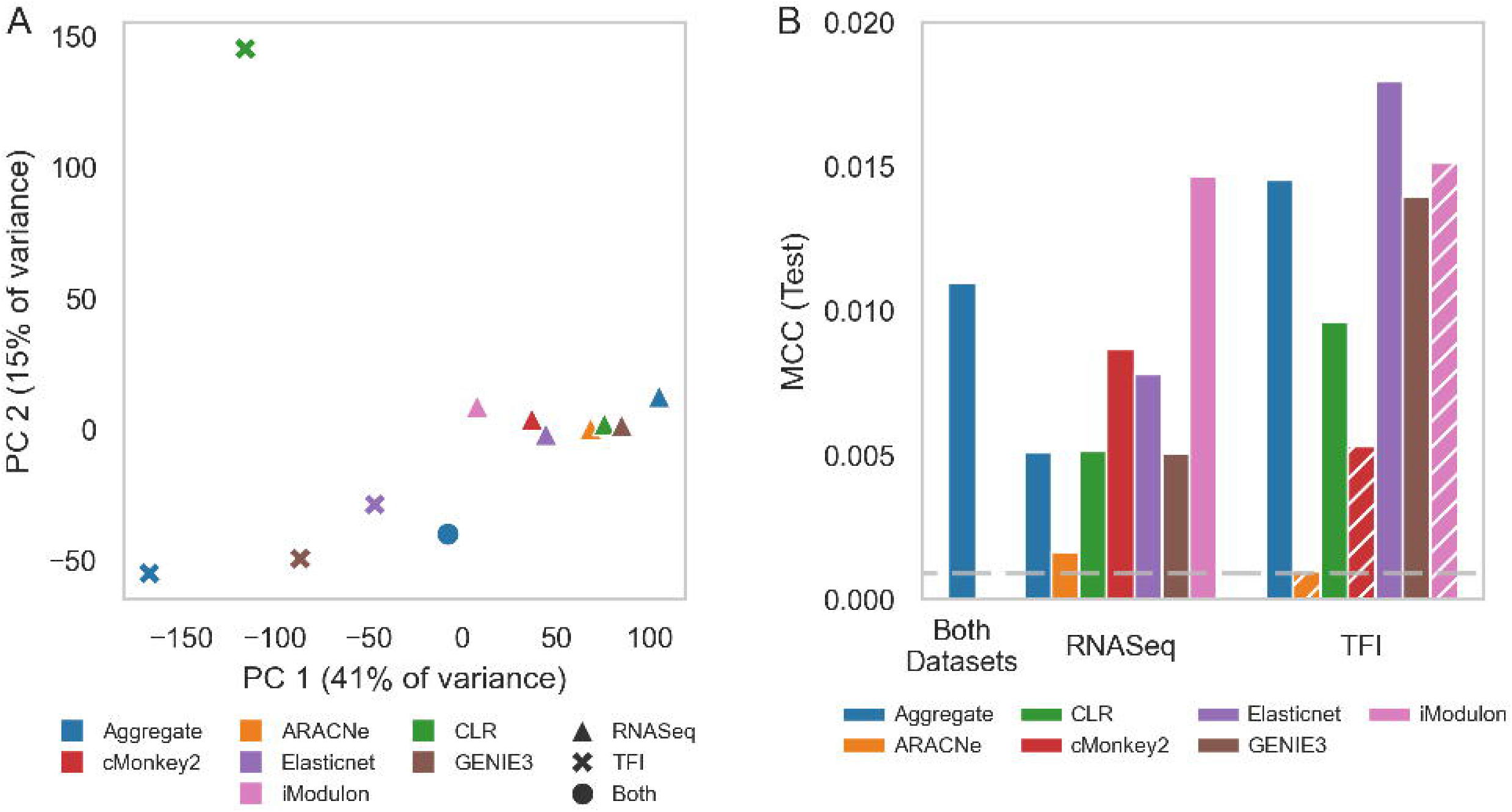
Overview of aggregate network. **(A)** PCA was performed on each of the generated networks. The networks inferred from the RNASeq compendium (triangle symbols) cluster to the right, whereas the networks inferred from the recombinant TF induction transcriptomes (x symbols) fall to the left. The dataset-level aggregates each cluster loosely with the same-dataset constituent networks at the horizontal extremes, whereas the overall aggregate falls near the centroid of all networks. **(B)** Performance of each inferred and aggregate network, calculated against a set of TF– target gene relationships defined by a ChIP-Seq DNA-binding investigation of recombinant TFI strains [3], as measured by Matthews correlation coefficient (MCC). MCC quantifies the level of correlation between the two sets, with higher values indicating more correspondence. Blue bars depict the MCC for aggregate networks; the other colors depict the MCC for the individual inferred networks. Hatched bars indicate networks that were excluded from aggregation. The horizontal dashed line represents the 95th percentile MCC performance of 1000 randomly generated networks. Note that the excluded iModulon/TF induction network scores relatively highly by this metric, likely because of its size (∼7k edges, versus an average of ∼180k). See Methods for information about the exclusion criteria.

We rank-aggregated the resulting 12 networks to consolidate regulatory relationships across the individual inference methods. The resulting aggregate network has 68,226 regulatory interactions that connect 214 transcriptional regulators with 4,027 target genes. Of these interactions, 37,236 are associated with transcriptional activation across conditions, 15,820 interactions are associated with transcriptional repression across conditions, 1,496 relationships are predicted to be either activating or repressing, depending on the environmental condition, and 11,766 regulatory relationships have an undetermined regulatory directionality (**Supplementary Table 4**). These interactions represent both direct, biophysical regulatory events as well as indirect regulatory relationships mediated by downstream regulators. These interactions also represent the union of regulatory relationships that are active in at least a subset of all the different environmental conditions profiled in our assembled source RNA-seq compendium and TF induction profiling datasets. Notably, not all these regulatory relationships will be active under all environmental conditions. The distribution of regulatory interactions per TF largely follows a power law distribution consistent with the scale free networks found to represent transcriptional regulation in other bacteria (**Supplementary** Figure 4). We found a deviation between the distribution of our aggregate network and the expected power law distribution for regulators with relatively few target genes. This is likely due to the inclusion of indirect regulatory relationships and relationships that are active under some but not all environmental conditions. The networks can be viewed at https://tfnetwork.streamlit.app/, and the TF-gene interactions are described in **Supplementary Table 4**.

To validate the connectivity of our aggregate network, we benchmarked it against experimentally profiled TF binding data we previously profiled by ChIP-seq in the TFI strains under log-phase broth culture [3]. To assemble a high-confidence regulatory association dataset, we included only significant ChIP-seq peaks associated with TF binding in the promoter region of target genes. We evaluated overlap between this high-confidence ChIP-seq regulatory interaction dataset and our inferred regulatory networks with the Matthews correlation coefficient (MCC). We find that most of the inferred networks that we generated had significant MCCs, and that the aggregate network outperforms the majority of inferred networks using individual methods (**Supplementary** Figure 3), whilst still retaining a large number of regulatory relationships (most of the better performing individual inference networks have relatively few regulatory interactions).

We also assessed the extent to which the regulatory relationships captured by our aggregate network preserved biological functional relationships between the regulating TFs and the target genes. For TFs with clear literature characterization of its function, we found a high degree of correspondence with the gene ontologies and annotated functions of its regulated target genes (**Table 1**, **Supplementary Table 5**). For example, Rv3574 (*kstR*) is a TF that has been linked to regulating cholesterol metabolism [54], and the target genes associated with *kstR* in our aggregate network also have gene ontology annotations linked to cholesterol metabolism (**Table 1**). Additionally, toxin-antitoxin target genes were enriched for growth regulation, highlighting that the regulatory relationships captured by the aggregate network include indirect regulatory relations. Collectively, this suggests the significant ontology and functional annotation enrichments made for genes and TFs that are currently poorly annotated represent testable hypotheses for function – this is one of the major advances from the aggregate network.

### 3.3 Network component analyses reveal per-sample Mtb TF activities under different conditions

Understanding when TFs are actively exerting their regulatory influence on their target genes can reveal mechanistic insights into bacterial physiology and stress response. Network component analysis (NCA) is an efficient way of estimating these TFA profiles from expression data by using a TRN to perform matrix decomposition [14]. Robust NCA (ROBNCA) is a variant of NCA that improves the performance of NCA calculations on noisy data with outlier measurements [43]. We applied ROBNCA to estimate TFAs corresponding to each sample in our TFI microarray and RNA-seq compendium.

To first determine and validate the ROBNCA TFA estimation approach on our data, we performed ROBNCA on the TFI microarray data using the aggregate network inferred only from the TFI data, as well as on 10 randomized networks to be used as negative controls. We hypothesized that if the estimated TFAs represent true TF activities, with high TFAs indicating strong net activator activity and low TFAs indicating strong net repressor activity, then the percentile ranks of TFAs for highly expressed TFs should be either very high or very low in their corresponding TFI strains. On the other hand, if the ROBNCA-calculated TFAs were spurious, then the TFA percentile ranks should be statistically indistinguishable from the TFA percentile ranks from randomized networks.

For each of the 208 TFI strains within the microarray expression dataset, we averaged the TFAs for all TFs across their biological replicates. We rank ordered TFs by their activities for each TFI strain, calculated the rank percentile activity of the induced TF for each TFI strain, and analyzed the distribution of these percentiles (**Figure 3A**). For the TFI microarray network, 31 TFs were ranked in the highest or lowest 15% of TFA ranks (greater than 1 standard deviation from the mean), implying that these TFs were the dominant regulators active in their respective TFI strain profiling condition. Interestingly, 91 TFs had TFAs in the middle 30% from 35-65%. These TFs were fairly uniformly distributed suggesting their related transcriptional programs were likely cross-regulated by other TFs.

**Figure 3:**
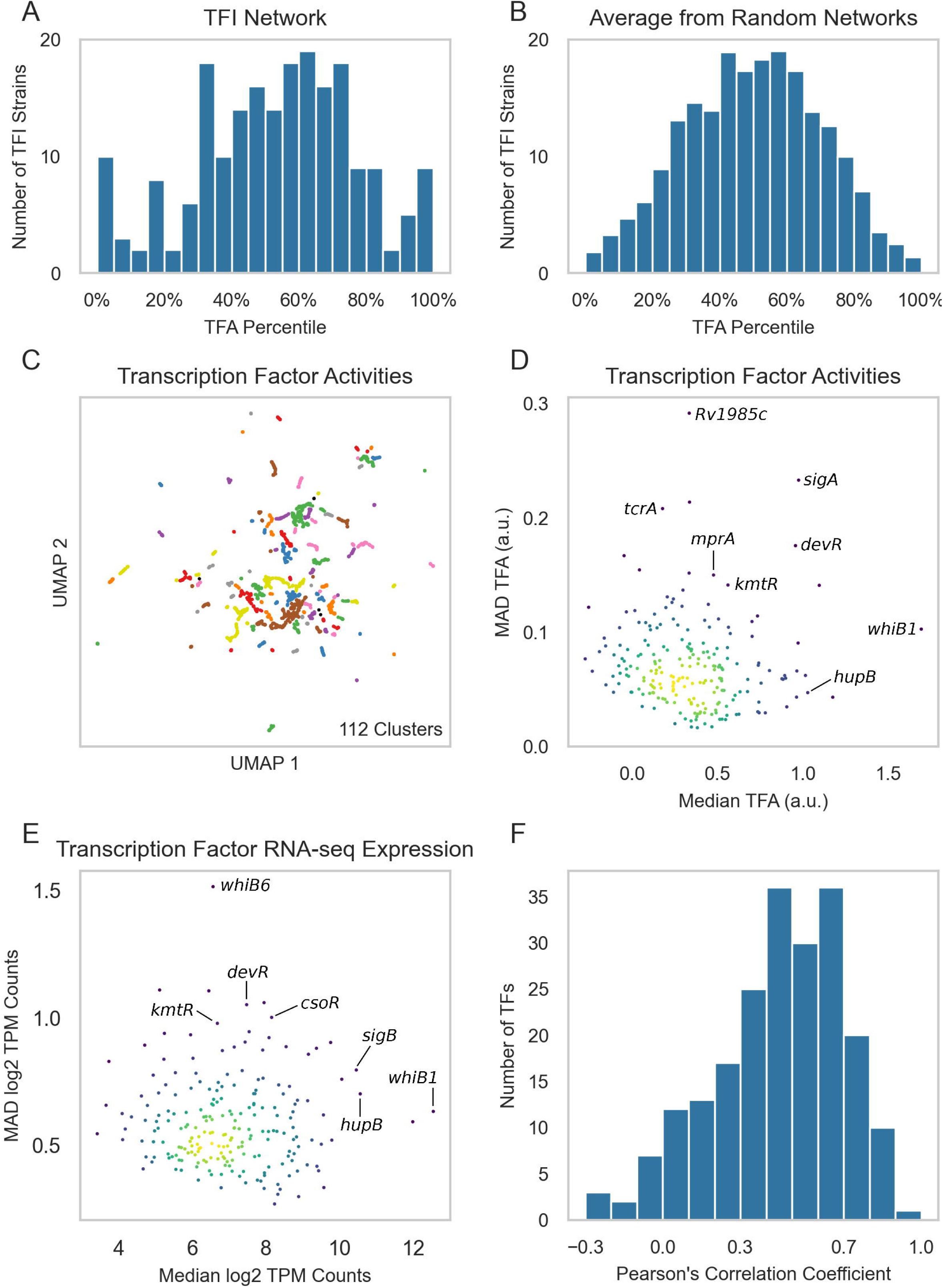
Compendium-wide transcription factor activities. (A) Distribution of the TFA rank percentiles for each induced TF in each strain from the TFI microarray dataset. ROBNCA was applied to the TFI microarray dataset using the network specifically inferred from the TFI dataset. For each sample, rank percentiles were computed for each TFA. TFAs were averaged across biological replicates for each TFI strain. Histogram depicts the percentile rank for TFAs corresponding to the over-expressed gene in each TFI strain. (B) Averaged distribution of TFA percentile ranks from ROBNCA using 10 randomized networks (**Supplementary** Figure 6). (C) UMAP visualization of samples from the normalized and batch corrected RNA-seq compendium as determined by TFA. UMAP and DBSCAN analyses reveal 112 clusters of samples with similar TFAs. (D) Median vs. MAD plot of activity for each TF across the RNA-seq compendium. (E) Median vs. MAD plot of expression for each TF across the RNA-seq compendium. (F) Distribution of Pearson’s correlation coefficients between expression and activity for each TF across the RNA-seq compendium.

Importantly, this suggested that induction of TF expression alone may be insufficient for fully inducing some transcriptional programs, thus supporting the use of TFAs over untransformed gene expression for downstream analysis.

We performed similar calculations for each of the randomized networks **(Supplementary** Figure 6) and averaged the TFA rank percentiles for all TFs from each randomized network (**Figure 3B**).

We found that there were significantly fewer TFAs in the highest or lowest 15% of TFA ranks in these randomized networks than the TFAs calculated from the TFI expression dataset (p = 1.66e-49, z-test [55]). Similarly, there were significantly more TFAs in the middle 30% (p = 1.66e-49, z-test [55]). These differences between the ROBNCA-calculated TFA percentile distributions between TFI and randomized networks indicated that the TFAs estimated by ROBNCA were not spurious and likely reported on true biological condition-specific activities.

We next applied ROBNCA to our RNA-seq compendium using the TRN inferred from the RNA-seq compendium. UMAP and DBSCAN analyses revealed that the level of biological diversity of ROBNCA-predicted TFAs was similar to the diversity within the expression compendium, with 112 clusters of TFAs across the 3,496 samples (versus 142 for untransformed expression; **Figure 3C**).

Amongst the TFs with the highest level of median activity were the essential nitric oxide-sensing Rv3219 (*whiB1*), histone-like protein Rv2986c (*hupB*), and sigma factor Rv2703 (*sigA*) (**Figure 3D**). Each of these would be expected to be constitutively active in live Mtb cells. Also consistent with expectation, the well-characterized stress response regulators Rv3133c (*devR*), Rv1994c (*cmtR*), Rv0827c (*kmtR*) and two-component system regulators Rv0602c (*tcrA*) and Rv0981 (*mprA*) were amongst the TFs with the highest TFA MAD.

Interestingly, the distribution of TFAs appeared different from the distribution of TF expression levels measured for each RNA-seq sample across the compendium (**Figure 3E**). We tested the correlation of expression level vs. activity for each TF across the entire compendium and found that expression and activity were only moderately correlated across the dataset (Pearson’s r = 0.48 ± 0.16 median ± MAD) (**Figure 3F**). 31 TFs were strongly correlated (|Pearson’s r| ≥ 0.7), 66 TFs were moderately correlated (0.7 > |r| ≥ 0.5), and 61 TFs were weakly correlated (0.5 > |r| ≥ 0.3). Relatedly, both median and MAD expression and activity were only weakly correlated across all TFs (median: r = 0.43; MAD: r = 0.32). These analyses further support our observation that TF expression level is not the sole determinant for TFAs for most TFs. Rather, expression and activity convey two distinct but complementary insights into transcriptional regulation, highlighting the importance of accounting for network interactions when investigating transcriptional regulation. In particular, we posit that TFs with weak correlation between expression and activity may require allosteric or other post-translational modification to trigger activation of transcriptional regulation. This hypothesis can be tested in future studies.

### 3.4 Transcription factor activity profiles can predict condition-specific bacterial fitness

Because transcriptional regulation plays important roles in coordinating Mtb growth adaptations under stress, we asked whether our regulatory network models could be used to predict fitness consequences of TF regulatory activities. To test this hypothesis, we utilized gradient boosting machine learning to construct an interpretable TFA regression model designed to predict the fitness of each TFI strain during log-phase culture based on each strain’s calculated TFA profiles. We trained this model using the TFAs computed by ROBNCA from the RNA-seq compendium, paired with TFI fitness measurements that we previously collected in a Transcriptional Regulator Induced Phenotype (TRIP) screen [16]. This TFA–fitness regression model was able to explain 87% of the observed variation of growth between the TFI strains in the TRIP screen (**Supplementary** Figure 7).

To determine if this TFA–fitness regression model could predict changes in Mtb fitness or growth from new data that were not used to train the model (e.g., under differing experimental conditions), we generated fitness predictions with our model using transcriptomes that we profiled from Mtb cells undergoing hypoxia and reaeration stress. From the TFA profiles calculated for cells exposed to hypoxia, the TFA–fitness regression model predicted a significant decrease in growth that persisted for each of the timepoints profiled under hypoxia (**Figure 4A**, **Supplementary** Figure 8). From the TFA profiles calculated for cells under reaeration, the model predicted a rebound in Mtb growth comparable to growth levels experimentally measured during log-phase culture. The kinetics of the shifts in growth predicted by the TFA–fitness regression model aligned well with the experimental measurements of Mtb bacteriostasis in hypoxia, followed by growth during reaeration (**Figure 4A**, **Supplementary** Figure 8). Importantly, the experimental growth data from the hypoxia-reaeration time course aligned better with the predictions from the TFA regression model than from an analogous regression model trained from TF expression data alone (**Supplementary** Figure 10).

**Figure 4:**
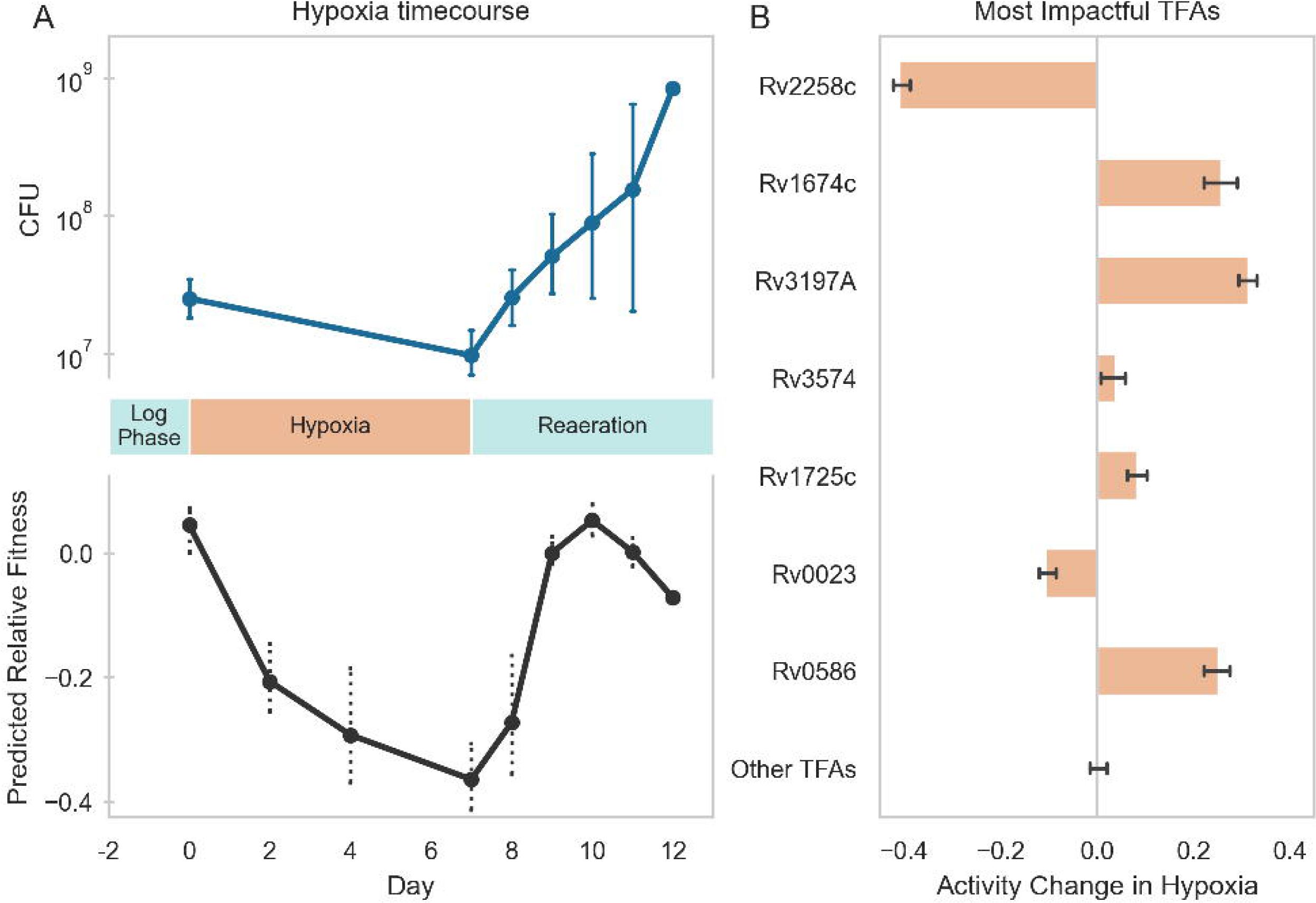
Machine learning model insights into Mtb growth through a hypoxic time-course. **(A)** *Top*: When Mtb grown for two days in log phase was subjected to hypoxic conditions (starting from day 0), the bacteria stopped growing for the duration of the imposed hypoxia, as indicated by the stable CFU between day 0 and day 7. When the culture was reintroduced to oxygen (“Reaeration”, starting from day 7), the bacteria resumed growth, as indicated by significantly higher CFU after day 8. *Bottom*: Our GBM model predicted a decrease in growth over the course of the period of hypoxia, and an increase in growth again upon reaeration, based only on transcriptional data measured over the course of the experiment. Each point represents an RNA-seq timepoint. **(B)** The GBM model can be interrogated to determine the primary drivers of the phenotype it predicts; when comparing the most impactful TFAs in hypoxic conditions (days 2-7) versus those in reliably reaerated conditions (days 9-12), 7 TFs were predicted to be particularly influential to the reduced growth in hypoxia versus reaeration, each contributing at least 5% of the total absolute impact predicted by the model. Shown here is the mean TFA change for each of the impactful TFs across days 2-7; other TFAs show no net activity change overall (see Methods for details on TFA change calculation).

These results further support our premise that TFAs more effectively capture condition-specific transcriptional regulation than TF expression alone and implies that the activation and regulation of transcriptional programs under hypoxia and reaeration may involve allosteric or other post-transcriptional mechanisms.

Because the TFA–fitness regression model is openly interpretable, we examined which TFAs most strongly predicted the fitness changes under hypoxia and reaeration. We found that our TFA– fitness regression model predicts that growth restriction during hypoxia is primarily driven by the activities of 7 TFs whose TFA profiles changed significantly during hypoxia (**Figure 4B**).

Importantly, each of these TFs have direct or indirect links to hypoxia in the literature (**Supplementary** Figure 9, **Supplementary Table 7**), thus further validating these model predictions and the use of TFAs as a lens into condition-specific stress response biology.

## 4. Discussion

Understanding the molecular drivers of phenotypic changes in an organism is a fundamental goal of biological research. In this study, we applied machine learning approaches to construct an interpretable TFA–fitness regression model that can utilize Mtb TRNs to predict experimentally measured changes in Mtb growth state in diverse environmental conditions. Our models build upon existing experimental profiling and network inference modeling efforts to characterize Mtb transcriptional regulation by integrating the data and algorithms developed in these prior studies [2; 3; 4; 5; 6; 7; 14; 15; 43]. Moreover, by integrating Mtb fitness profiling data from TRIP, our models have also enabled direct prediction of growth/survival phenotypic outcomes from condition-specific gene expression data inputs.

Our “wisdom of crowds” approach for inferring transcriptional regulatory interactions yielded significant enrichment of known regulatory relationships while also expanding the scope of represented experimental conditions. Our resulting TRN is substantially larger than the networks inferred by individual algorithms, while enriched for experimentally validated interactions. This highlights the utility of ensemble inference algorithms, as has been previously shown for regulatory network inference in other bacteria [15].

Importantly, our results demonstrate how network models can generate hypotheses on gene function in at least two complementary ways. First, we show by gene ontology enrichment analysis that there is significant correlation between the annotated function of a TF’s target genes and the condition-specific regulatory function of the TF. It is important to note that the regulatory interactions identified by our aggregate TRN includes both direct regulatory interactions involving physical interactions between a TF and its target gene as well as indirect associations mediated by other factors. Both direct and indirect regulatory associations are important for coordinating changes in bacterial physiology [56], so it is expected that both types of interactions share annotated ontologies. Because ∼25% of Mtb genes lack functional annotation [57], we think the regulatory relationships identified in our TRN can aid basic microbiological efforts in investigating Mtb gene function by generating hypotheses for the functions of these poorly characterized or unknown genes (**Supplementary Table 5)**.

Second, we show that TFA regression models can be trained to link condition-specific TFAs with TF fitness in log-phase broth culture to predict Mtb fitness under stress. Notably, we show that our TFA regression model was able to predict Mtb growth and bacteriostasis under hypoxia and reaeration – environmental conditions not used in training the TFA regression model. Our results biologically suggest that TFAs are a useful determinant of condition-specific changes in bacterial growth, and that the estimated TFA is more predictive of growth phenotypes that TF expression alone. This is consistent with expectation as Mtb uses transcriptional regulation to orchestrate behavioral adaptations to varying environments, including in growth phenotypes. Our modeling also enables inspection of which TFAs are driving the predicted bacterial fitness outcomes. This can inform the generation of hypotheses on the mechanisms underlying how TFs and their corresponding transcriptional programs are activated (e.g., via allosteric mechanisms and/or network interactions). Our TRN and TFA–fitness models could potentially inform the identification of regulatory mechanisms mediating Mtb response and adaptation to other clinically relevant stress conditions where gene expression profiling data are available. The TFs and target genes highlighted by these models may potentially represent future intervention targets aimed at modulating Mtb fitness in a therapeutically beneficial way. In light of the growing crisis of antimicrobial resistance [58] and multi- and extensively-drug-resistant tuberculosis [59], we think our approach will be important for curing tuberculosis disease [60].

More broadly, our work here demonstrates how network models can be utilized for biologically meaningful interpretable machine learning applications. A fundamental challenge in current machine learning activities is the difficulty in understanding how a trained machine learning model makes predictions [61; 62]. We previously demonstrated that machine learning regression models can be used to elucidate metabolic mechanisms underlying antibiotic lethality in *E. coli* [63], as well as to predict multidrug interaction outcomes in Mtb [50]. Our study here analogously extends this approach by training a regression model on TFAs estimated from TRN analyses to predict changes in Mtb growth state. The advantage of this strategy over other contemporary machine learning approaches is the direct utilization of prior knowledge encompassed by biological network models, which directly enable the generation of hypotheses for mechanisms linking network interactions to cell phenotypes. These hypotheses can then be experimentally tested [50; 63] and used as the basis for further mechanistic study [64] and investigation of translational potential.

Looking forward, we envision that this approach and our TFA regression model can be useful for several facets of tuberculosis research. We demonstrated that our model can be used to predict changes in Mtb growth state under environmental stress, which may inform the design of growth state assays under conditions where standard microbiological tools are not feasible. There is increasing appreciation that Mtb drug susceptibility is regulated by its environment [65; 66]. Our TFA–fitness regression model can be used to elucidate the molecular mechanisms underlying these phenotypes. Moreover, functional genetic datasets are becoming increasingly available using different technologies [16; 67; 68; 69; 70; 71; 72]. These data can be applied to train next-generation TFA–fitness regression models with improved predictive power. Finally, detailed characterizations of Mtb clinical strains are now providing significant insights into the how mutations or other forms of genomic diversity regulate drug susceptibility in human patients [72; 73; 74; 75]. We envision the TRN and TFA-fitness regression framework established here can be extended not only to study the mechanistic basis for differences between drug susceptibility amongst clinical isolates, but also to anticipate the drug susceptibility of new clinical isolates as they become curated.

## Author Contributions

E.B.: Formal Analysis, Investigation, Methodology, Software, Validation, Visualization, Writing – original draft, Writing – review, editing; E.P.: Data curation, Formal analysis, Visualization, Writing – review, editing; O.G.: Formal Analysis, Investigation, Methodology, Software, Validation, Visualization, Writing – review, editing; B.T.G.: Data curation, Software, Visualization, Writing – review, editing; T.R.R.: Investigation, Resources, Methodology, Writing – review, editing; D.R.S.: Investigation, Resources, Methodology, Funding acquisition, Supervision, Writing – review, editing; J.H.Y.: Conceptualization, Funding acquisition, Investigation, Methodology, Project administration, Resources, Supervision, Visualization, Validation, Formal Analysis, Writing – original draft, Writing – review, editing; S.M.: Conceptualization, Funding acquisition, Investigation, Methodology, Project administration, Resources, Supervision, Visualization, Formal Analysis, Writing – original draft, Writing – review, editing.

## Acknowledgments

We thank Research Scientific Computing at Seattle Children’s Research Institute for providing HPC resources that have contributed to this investigation. We thank Avi Shah for helpful discussions, and Jessica Assadi and Robert Morrison for technical assistance.

## Supplementary Material

1. **Supplementary Figure 1 UMAP.** Hyperparameter optimization was performed on UMAPs from the (A) TFI microarray compendium, (B) RNA-seq compendium, or (C) TFAs calculated from the RNA-seq compendium. ε was varied from 0.1 to 10 on a logarithmic scale and numbers of clusters (left), numbers of outliers (center), and maximum cluster size (right) were computed for each ε. ε was selected from the elbow of the outliers plot (ε = 0.281 for TFI data, 0.309 for RNA-seq compendium and estimated TFAs).
2. **Supplementary Figure 2 Inferred network validation.** Distribution of the ranks, in each network, of edges shared with the validation dataset from Sanz et al., 2011, [8] from each network. Each histogram is divided into 32 bins. Horizontal dashed lines represent the expected number of random matches between each network and the validation dataset. Truncation was performed on these networks at the first bin where the count dropped below the dashed line (see **Methods**). Panels with hashed backgrounds (B, F, and L) represent networks that were excluded from the aggregation due to insufficient enrichment.
3. **Supplementary Figure 3. Inferred network performance.** Performance of each inferred and aggregate network, calculated against a set of TF–target gene relationships identified by Sanz et al., 2011 [8] (see **Methods**), as measured by Matthews correlation coefficient (MCC). MCC quantifies the level of correlation between the two independent sets of relationships. Higher values indicate greater correlation. The blue bars depict the MCC for the dataset-level and overall aggregates. Other colors are used to depict the MCC for the individually inferred networks. Hatched bars indicate the networks that were excluded from aggregation. The horizontal dashed line represents the 95th percentile MCC performance of 1,000 randomly generated networks. See Methods for exclusion criteria.
4. **Supplementary Figure 4 TRN properties.** Out-degree distribution of TF-gene interactions (edges) from the overall aggregate network. This distribution significantly differs from a power law distribution on the left side of the plot, likely because the network includes indirect interactions. These will deflate counts of low-degree TFs (nodes) and inflate counts of higher- degree nodes.
5. **Supplementary Figure 5 Assignment of activating vs repressing regulatory interactions.** Flow chart depicting the logic used to assign directionality to regulatory relationships. Abbreviations used are defined in the legend in the bottom left.
6. **Supplementary Figure 6 TFA rank percentiles for randomized networks.** TRNs were randomized 10 times. For each random network, ROBNCA was used to compute TFAs for the TFI dataset. Rank percentiles were assigned to each TFA for each TFI microarray profile and averaged across replicates for each TFI strain. Plotted are TFA rank percentile distributions for all over-expressed TFs corresponding to their respective TFI strain from each randomized network.
7. **Supplementary Figure 7. TFA-fitness regression model performance.** (A) Fitness values predicted by the gradient boosted machine (GBM) model versus the experimentally measured values supplied to the model upon training. The line of best fit depicts the relationship between predicted and measured values. The slope of this line is slightly less than 1, indicating that the regression model modestly underestimates relative fitness changes. The model achieved a coefficient of determination (R^2^) of 0.87 against its training set, indicating that the model can explain 87% of the variation in fitness from the TRIP screen. (B) Residuals of the model predictions versus measured values form a roughly normal distribution, indicating a lack of bias and overall reliable predictive ability.
8. **Supplementary Figure 8 TFA hypoxia prediction.** Our TFA-fitness regression model predicted a decrease in growth over the course of the period of hypoxia, and an increase in growth again upon reaeration, based only on transcriptional data measured over the course of the experiment (each point represents an RNA-seq timepoint), in both the empty plasmid strain (blue) and wild-type H37Rv (orange).
9. **Supplementary Figure 9 Hypoxia-responsive TFAs.** The TFA-fitness regression model can be interrogated to determine drivers of hypoxia by comparing the most impactful TFAs under hypoxia (days 2-7) versus reaeration (days 9-12). 7 TFs were most important for predicting reduced growth under hypoxia versus reaeration. Each contributes at least 5% to total model predictions. Depicted is the mean change in TFA for each of the impactful TFs across days 2-7 (orange) versus days 9-12 (cyan). Other TFAs show negligible changes in activity across hypoxia or (see Methods for details on calculations for changes in TFA).
10. **Supplementary Figure 10 TF expression hypoxia prediction.** Hypoxia and reaeration fitness changes predicted by a GBM model trained using only TF expression data instead of TFAs.
11. **Supplementary Table 1 Expression data from the TFI microarray dataset.** Batch correction group assignments for each sample in the TFI microarray dataset. Smooth quantile normalized and microarray expression for all genes and all samples in the TFI microarray dataset. Median and MAD expression for each gene. Group assignments were used by the PySNAIL smooth quantile normalization algorithm for batch correction [18].
12. **Supplementary Table 2 Expression data from the RNA-seq expression compendium.** Batch correction group assignments for each sample in the RNA-seq compendium. Group assignments were used by the PySNAIL smooth quantile normalization algorithm for batch correction [18]. Median and MAD expression for each gene.
13. **Supplementary Table 3 Network inference methods.** Description of transcriptional regulatory network inference methods.
14. **Supplementary Table 4. Aggregate network directionality of regulation.** Summary of the assignments of activating (up) vs. repressing (down) regulatory interactions for all TF-gene regulatory interactions in the aggregate transcriptional regulatory network (TRN).
15. **Supplementary Table 5 TF Gene Ontology assignments.** GO enrichment for each transcriptional program regulated by each TF inferred by our aggregate TRN. (A) Annotated functions and a summary of GO enrichments found for targets from selected TFs. All TFs with at least 3 significant GO enrichment terms and a non-locus gene name in Mycobrowser [40]. 45 TFs meet these criteria. These data validate the accuracy of our network, as one would expect an accurate regulatory network to have target sets significantly enriched for the known functions of each TF. (B) Remaining TFs with at least 3 significant GO enrichments assigned by our analysis but without an annotated gene name (36 additional TFs). These data represent predictions for potentially novel TF functions. (C) All GO enrichments identified by our analysis were corrected for FDR with a cutoff of 0.05.
16. **Supplementary Table 6 Transcription Factor Activities.** Median and MAD expression and activity for each TF in the RNA-seq compendium. Pearson correlation coefficient between TF expression and TFA for each TF across all samples in the RNA-seq compendium.
17. **Supplementary Table 7** Overview of the top 7 most important TFAs for predicting fitness under hypoxia as identified by our TFA regression model, validated by published evidence for mechanistic activation under hypoxia [3; 6; 76; 77; 78; 79; 80; 81; 82; 83; 84; 85; 86].

## Data Availability Statement

The transcriptome datasets analyzed for this study can be found in the supplemental material and at https://tfnetwork.streamlit.app. The code and software implementations associated with this study can be found at https://github.com/Ma-Lab-Seattle-Childrens-CGIDR/Mtb-TFA-fitness-regression and https://hub.docker.com/repositories/malabcgidr.

