## Supplemental Data 1 for "Predicting bacterial fitness in Mycobacterium tuberculosis with transcriptional regulatory network-informed interpretable machine learning"

### *Supplementary Material*

#### 1 Supplementary Figures and Tables

##### 1.1 Supplementary Figures

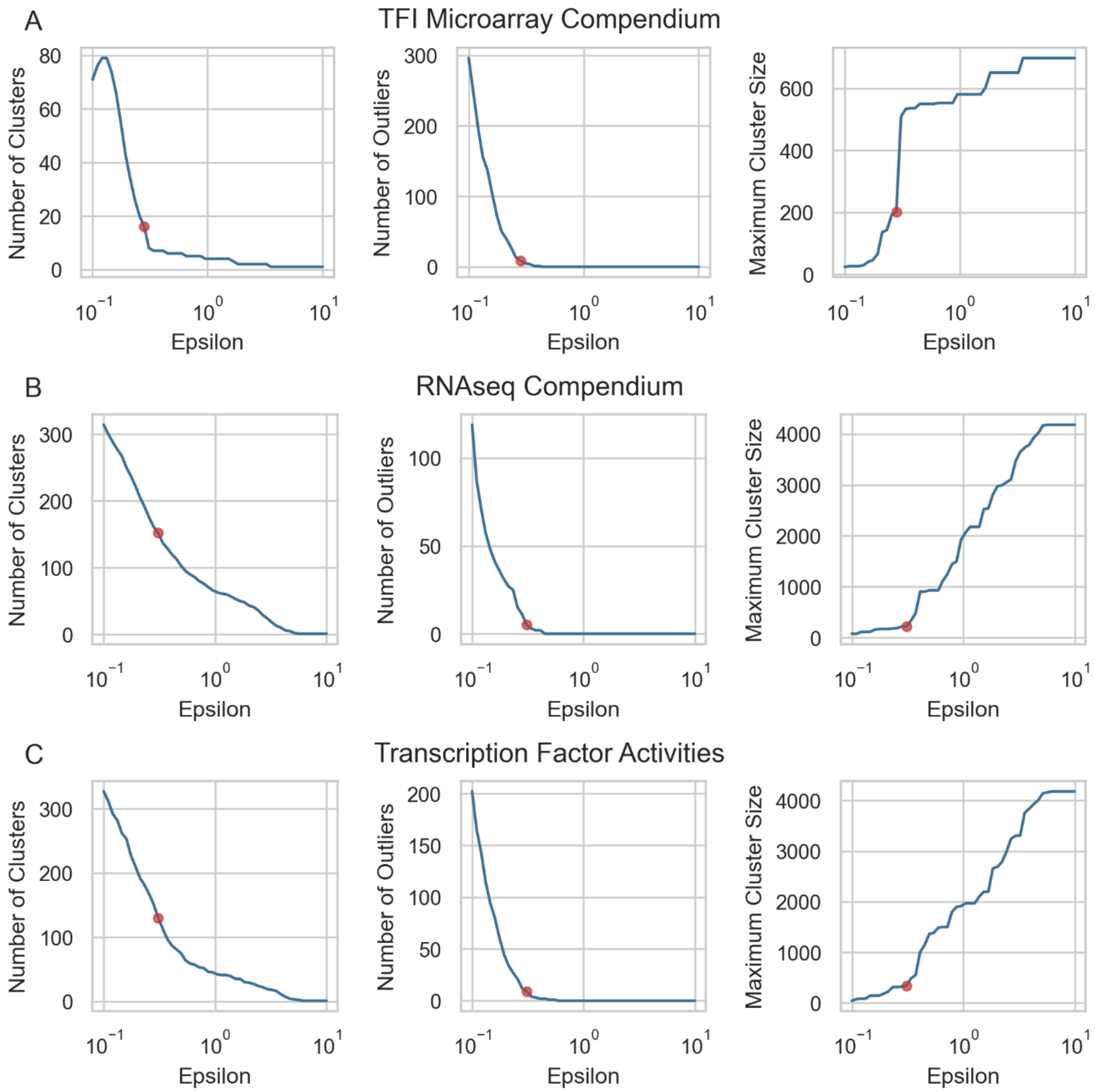


**Supplementary Figure 1 DBSCAN hyperparameter optimization.** Hyperparameter optimization was performed on UMAPs from the (A) TFI microarray compendium, (B) RNA-seq compendium, or (C) TFAs calculated from the RNA-seq compendium. $\varepsilon$ was varied from 0.1 to 10 on a logarithmic scale and numbers of clusters (left), numbers of outliers (center), and maximum cluster size (right) were computed for each $\varepsilon$. $\varepsilon$ was selected from the elbow of the outliers plot ($\varepsilon$ = 0.281 for TFI data, 0.309 for RNA-seq compendium and estimated TFAs).

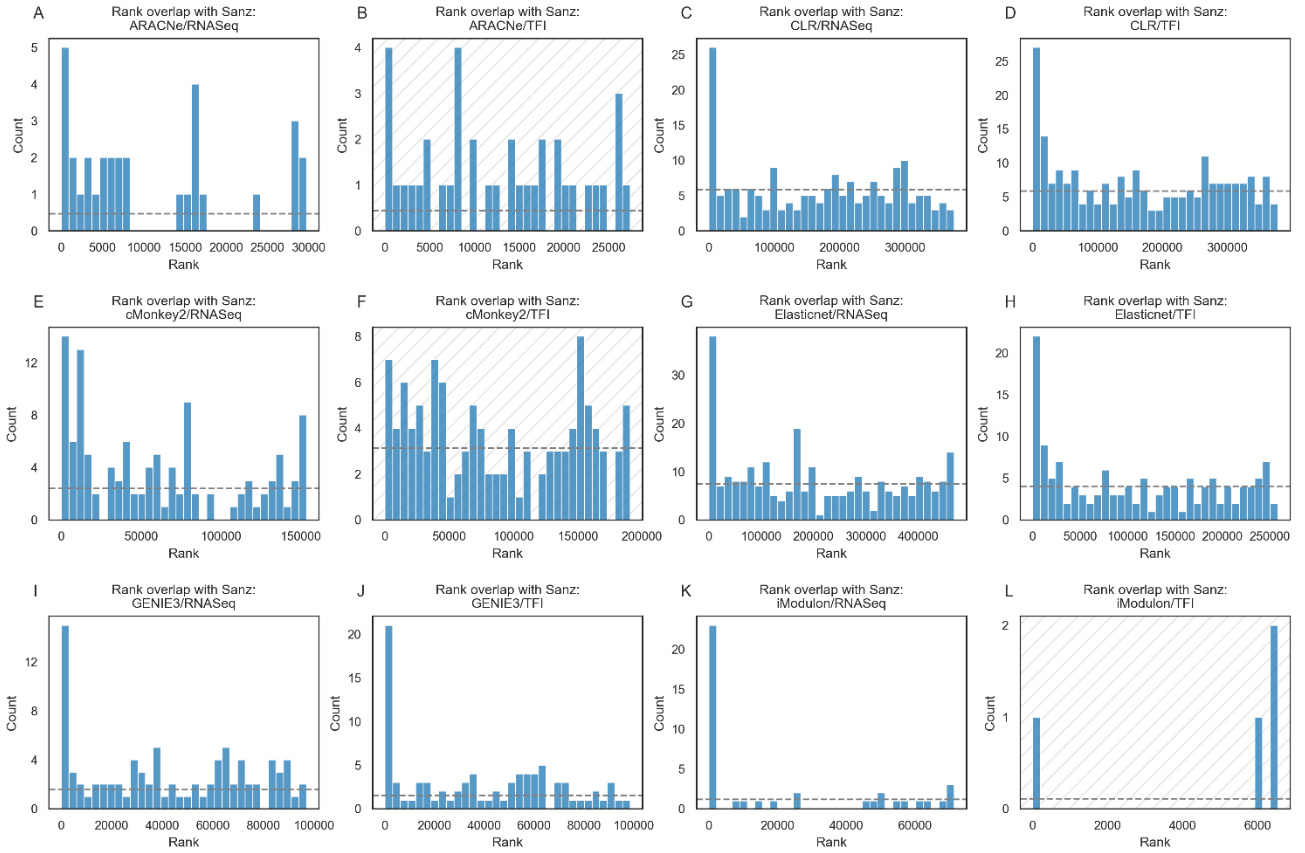


**Supplementary Figure 2 Inferred network validation.** Distribution of ranks for edges shared with the validation dataset from Sanz et al., 2011, [8] from each network. Each histogram is divided into 32 bins. Horizontal dashed lines represent the expected number of matches between each network and the validation dataset. Truncation was performed on these networks at the first bin where the count dropped below the dashed line (see Methods). Panels with hashed backgrounds (B, F, and L) represent networks that were excluded from the aggregation due to insufficient enrichment.

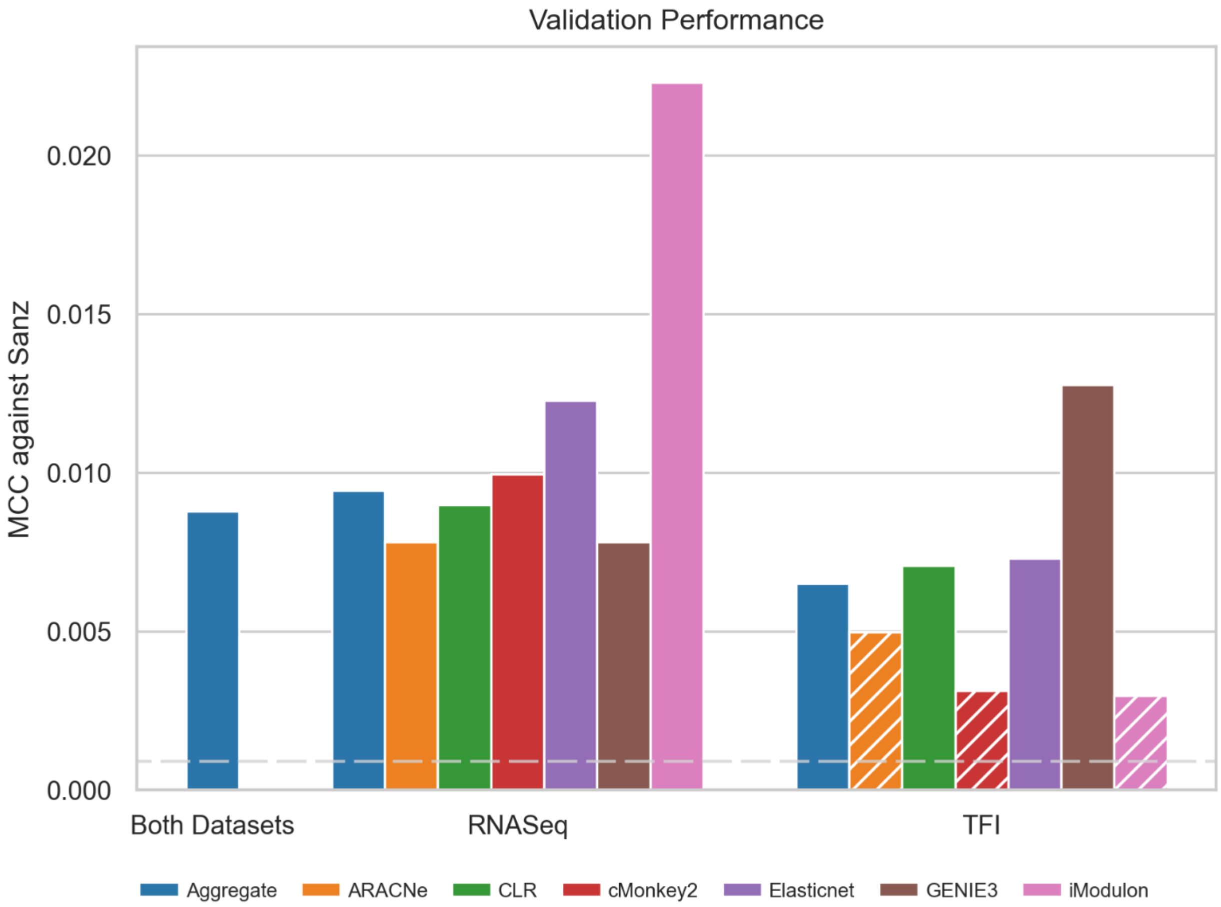


**Supplementary Figure 3 Inferred network performance.** Performance of each inferred and aggregate network, calculated against a set of TF–target gene relationships identified by Sanz et al., 2011 [8] (see Methods), as reported by the Matthews correlation coefficient (MCC). MCC quantifies the level of correlation between two independent sets of relationships. Higher values indicate greater correlation. The blue bars depict the MCC for the dataset-level and overall aggregates. Other colors are used to depict the MCC for individually inferred networks. Hatched bars indicate the networks that were excluded from aggregation. The horizontal dashed line represents the 95th percentile MCC performance for 1,000 randomly generated networks. See Methods for exclusion criteria.

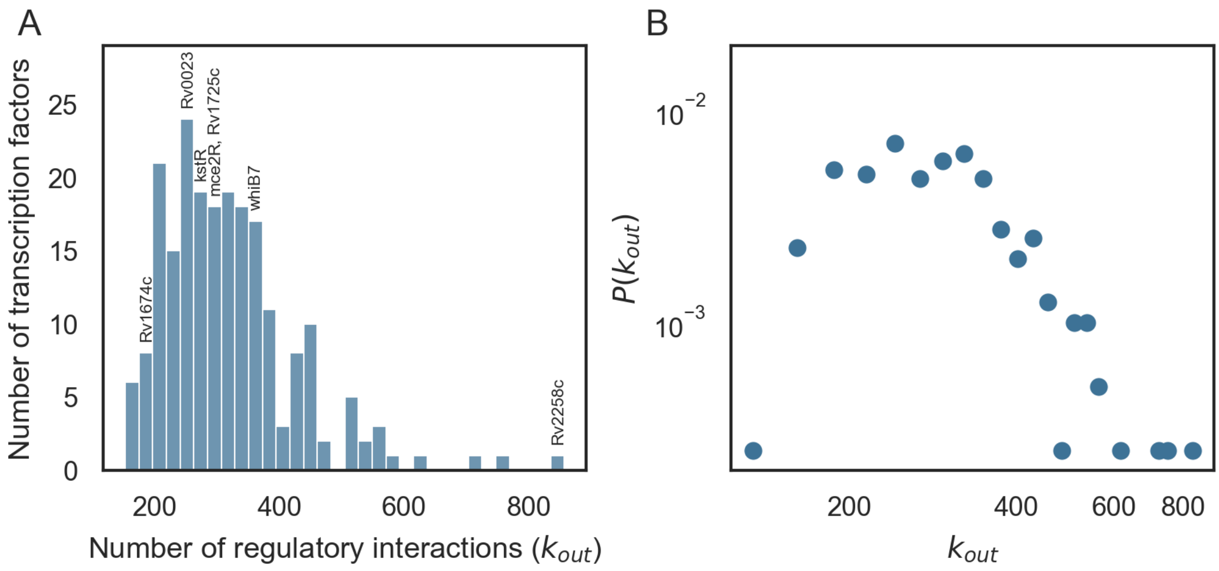


**Supplementary Figure 4 TRN properties.** Out-degree distribution of TF-gene interactions (edges) from the overall aggregate network. This distribution significantly differs from a power law distribution – likely because the network includes indirect interactions. These will deflate counts of low-degree TFs (nodes) and inflate counts of high degree nodes.

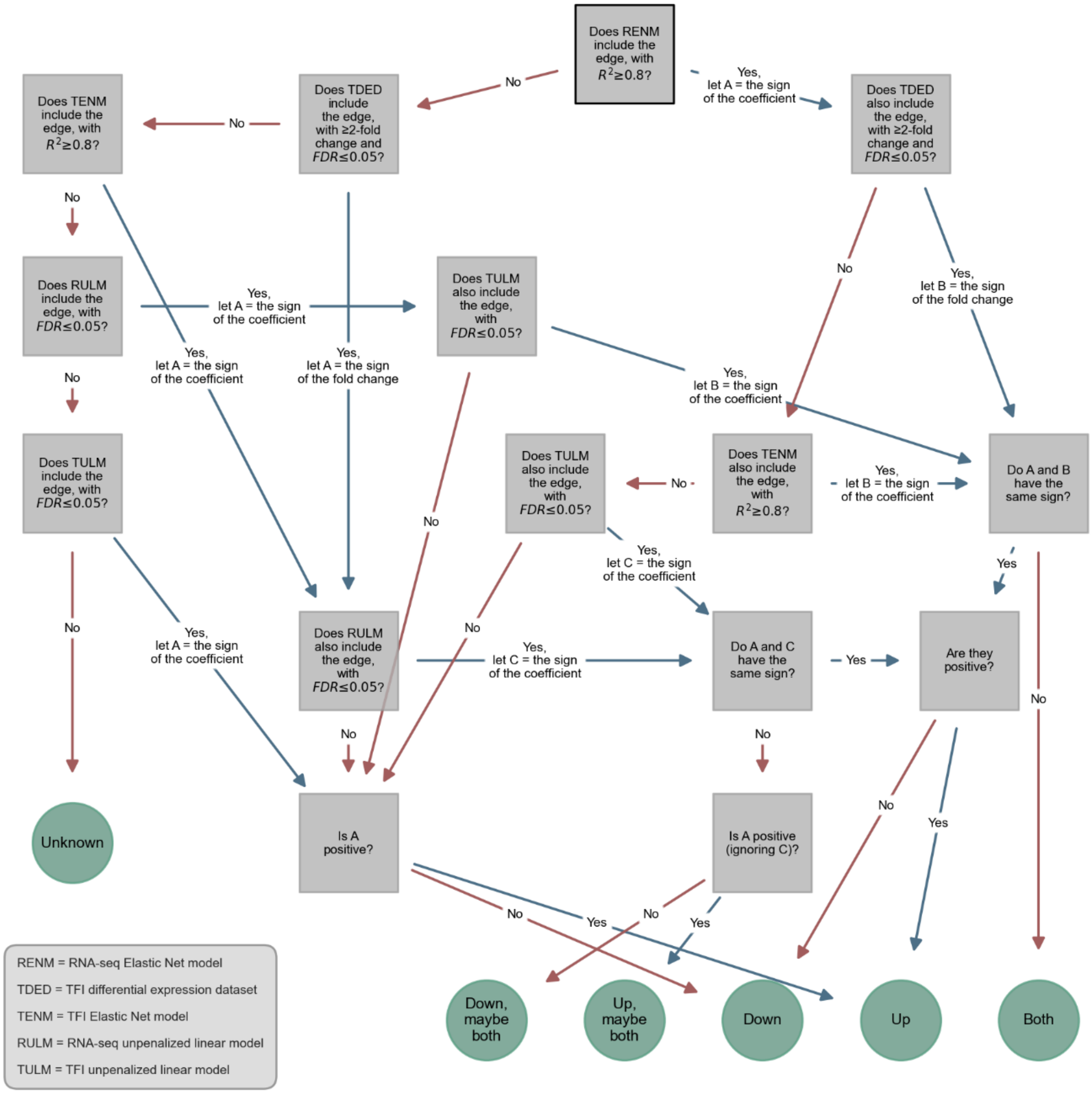


**Supplementary Figure 5 Assignment of activating vs repressing regulatory interactions.** Flow chart depicting the logic used to assign directionality to regulatory relationships. Abbreviations are defined in the legend in the bottom left.

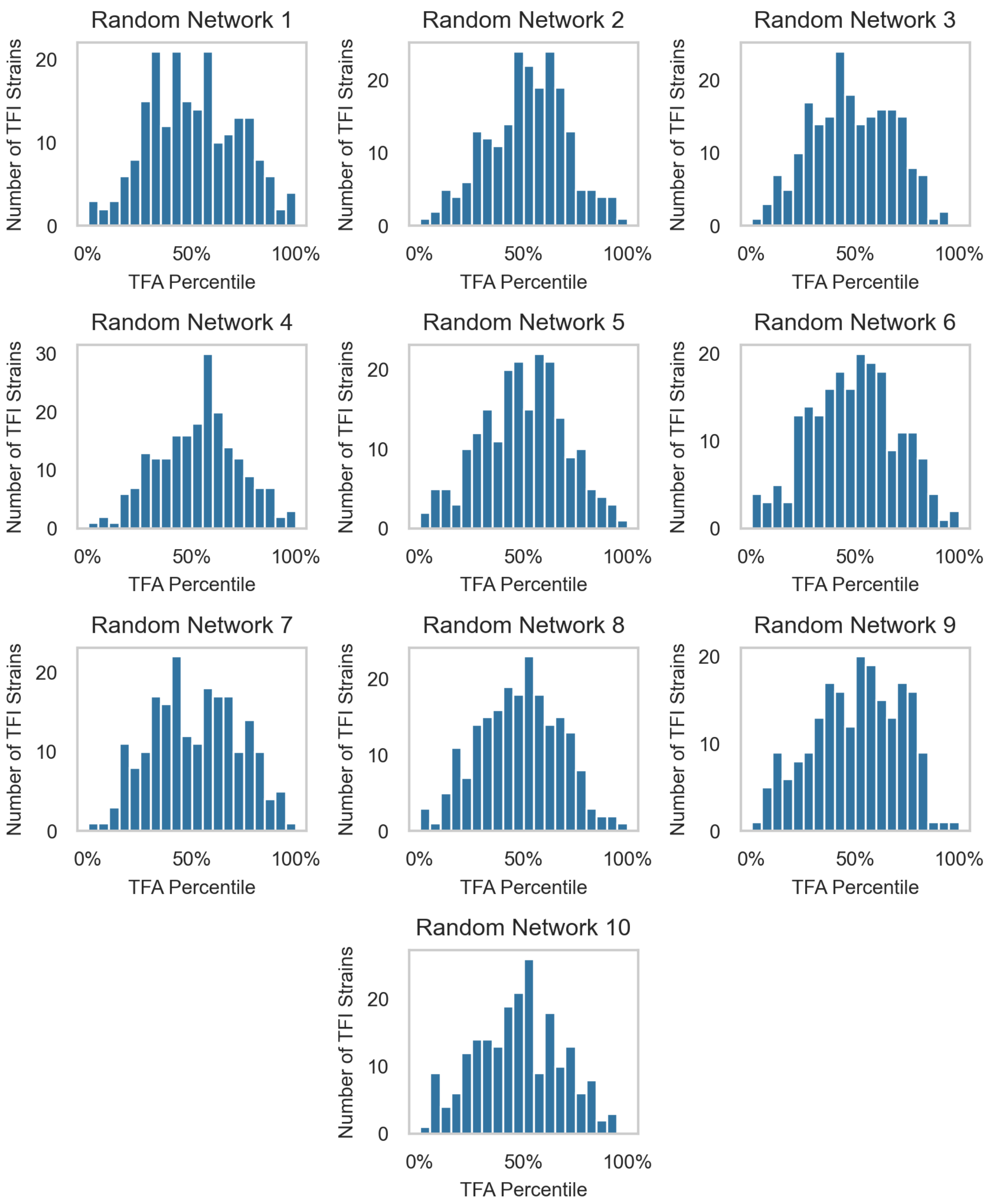


**Supplementary Figure 6 TFA rank percentiles for randomized networks.** TRNs were randomized 10 times. For each random network, ROBNCA was used to compute TFAs for the TFI dataset. Rank percentiles were assigned to each TFA for each TFI microarray profile and averaged across replicates for each TFI strain. Plotted are TFA rank percentile distributions for all over-expressed TFs corresponding to their respective TFI strain from each randomized network.

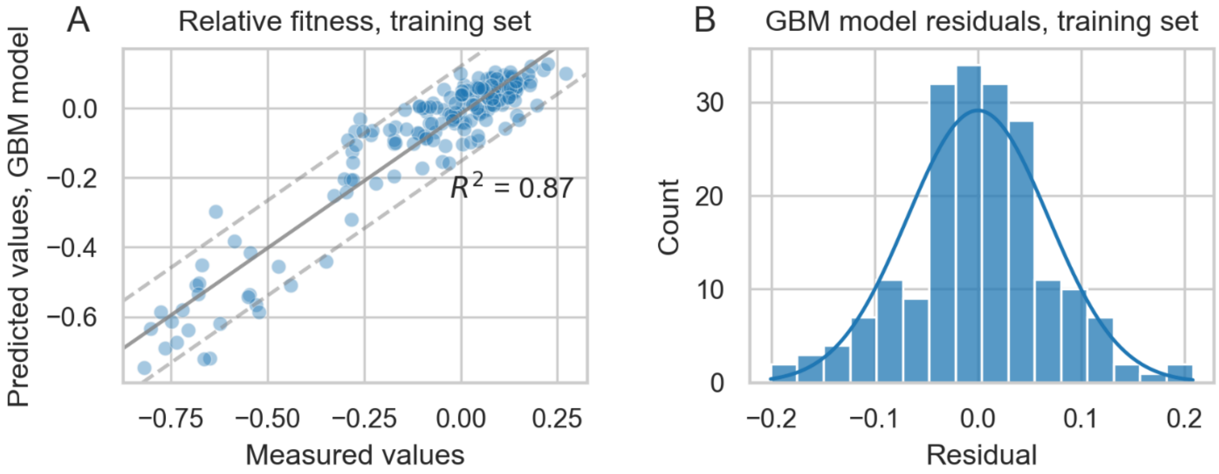


**Supplementary Figure 7 TFA-fitness regression model performance.** (A) Fitness values predicted by the gradient boosted machine (GBM) model versus the experimentally measured values supplied to the model upon training. The line of best fit depicts the relationship between predicted and measured values. The slope of this line is slightly less than 1, indicating that the regression model modestly underestimates relative fitness changes. The model achieved a coefficient of determination ($R^2$) of 0.87 against its training set, indicating that the model can explain 87% of the variation in fitness from the TRIP screen. (B) Residuals of the model predictions versus measured values form a roughly normal distribution, indicating a lack of bias and overall reliable predictive ability.

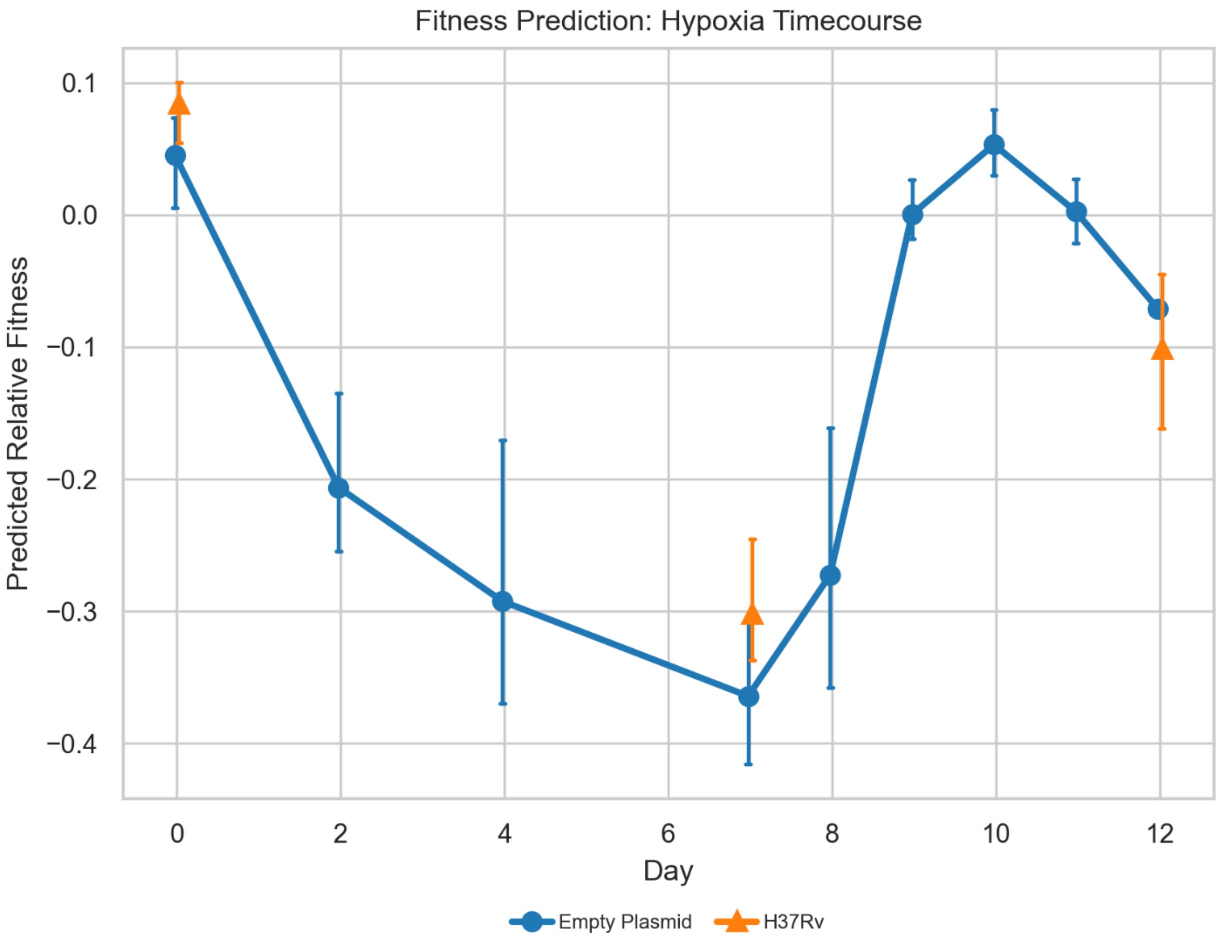


**Supplementary Figure 8 TFA hypoxia prediction.** Our TFA-fitness regression model predicted a decrease in growth over the course of the period of hypoxia, and an increase in growth again upon reaeration, based only on transcriptional data measured over the course of the experiment (each point represents an RNA-seq timepoint), in both the empty plasmid strain (blue) and wild-type H37Rv (orange).

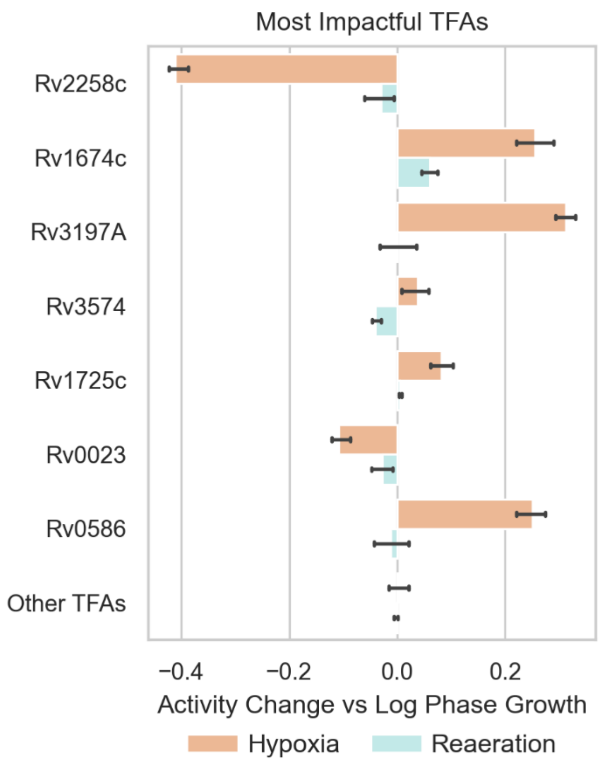


**Supplementary Figure 9 Hypoxia-responsive TFAs.** The TFA-fitness regression model can be interrogated to determine drivers of hypoxia by comparing the most impactful TFAs under hypoxia (days 2-7) versus reaeration (days 9-12). 7 TFs were most important for predicting reduced growth under hypoxia versus reaeration. Each contributes at least 5% to total model predictions. Depicted is the mean change in TFA for each of the impactful TFs across days 2-7 (orange) versus days 9-12 (cyan). Other TFAs show negligible changes in activity across hypoxia or (see Methods for details on calculations for changes in TFA).

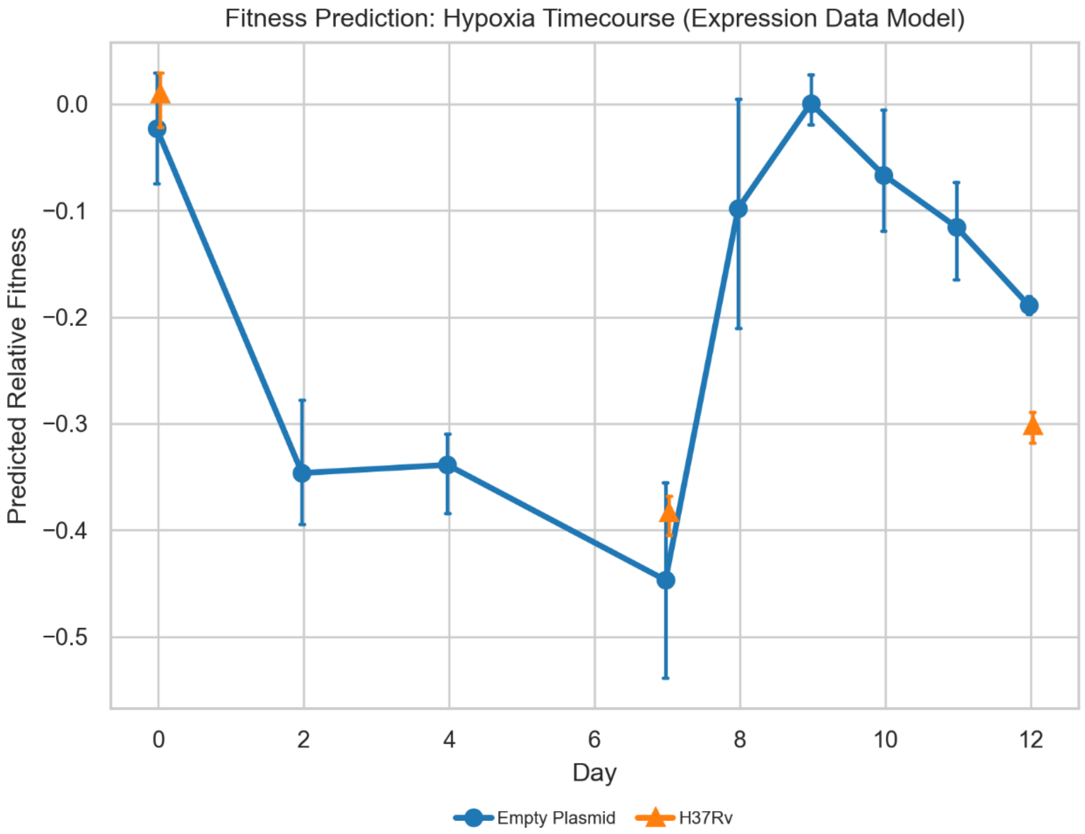


**Supplementary Figure 10 TF expression hypoxia prediction.** Hypoxia and reaeration fitness changes predicted by a GBM model trained using only TF expression data instead of TFAs.

#### 1.2 Supplementary Tables

**Supplementary Table 1 Expression data from the TFI microarray dataset.** Batch correction group assignments for each sample in the TFI microarray dataset. Smooth quantile normalized and microarray expression for all genes and all samples in the TFI microarray dataset. Median and MAD expression for each gene. Group assignments were used by the PySNAIL smooth quantile normalization algorithm for batch correction [18].
